# A novel membrane contact site in vestibular hair cells

**DOI:** 10.1101/2025.03.21.644540

**Authors:** Nicolas Brouilly, Nathalie Pujol, Fabrice Richard, Chantal Cazevieille, Jennifer Stone, Ruth Anne Eatock, Rémy Pujol

**Affiliations:** IBDM, Institut de Biologie du Développement de Marseille, Aix Marseille Univ, Marseille, France; CIML, Aix Marseille Univ, INSERM, CNRS, Turing Centre for Living Systems, Marseille, France; INM, Institut des Neurosciences de Montpellier, INSERM-Université de Montpellier, Montpellier, France; Department of Otolaryngology/Head and Neck Surgery, Virginia Merrill Bloedel Hearing Research Center and the Department of Otolaryngology-Head and Neck Surgery, University of Washington, Seattle WA, USA; Department of Neurobiology, University of Chicago, Chicago IL, USA

## Abstract

The mammalian vestibular system has two types of sensory receptor hair cells (HCs), each with different neurotransmission mechanisms. Type II HCs use ribbon synapses to transmit neurotransmitters like glutamate to afferent neurons. On the other hand, type I HCs are nearly engulfed by a calyceal afferent ending and also form ribbon synapses. These HCs regulate afferent activity through non-quantal transmission (NQT), which is faster than classic neurotransmitter release and may play a key role in stabilizing vision and balance during rapid head movements. Here, we describe a novel striated contact, present between the mouse type I HC and its calyceal afferent ending, and intimately associated with atypical plasma membrane-apposed (PMA) mitochondria. This distinctive arrangement has the potential to serve or modulate NQT.

---

The mammalian vestibular inner ear is in charge of body balance, sensing head motion and coordinating body movements. In the vestibule of amniotes, there are two types of sensory receptor cells, called hair cells (HCs), which differ dramatically in their mode of neurotransmission to primary afferent neurons. The afferent synapse of type II HCs, like that at most hair cells, is a ribbon synapse, with glutamate vesicles arrayed around a presynaptic ‘ribbon’ and facing a postsynaptic afferent bouton. In contrast, the type I HC is nearly fully engulfed by a calyceal afferent ending (Figure 1 A,B). The type I HC also forms ribbon synapses with the calyx (Lysakowski, 1999; Michanski et al., 2023), and it also regulates vestibular afferent activity via non-quantal transmission (NQT) (Eatock, 2018; Govindaraju et al., 2023; Songer and Eatock, 2013). This NQT is faster than classic quantal transmission (Songer and Eatock, 2013) and may help explain the remarkable ability of the vestibular system to stabilise vision and balance during rapid head motions (Govindaraju et al., 2023). Here, we describe a novel striated contact, present between the mouse type I HC and its calyceal afferent ending, and intimately associated with atypical plasma membrane-apposed (PMA) mitochondria. This distinctive arrangement has the potential to serve or modulate NQT.

**Figure 1:**
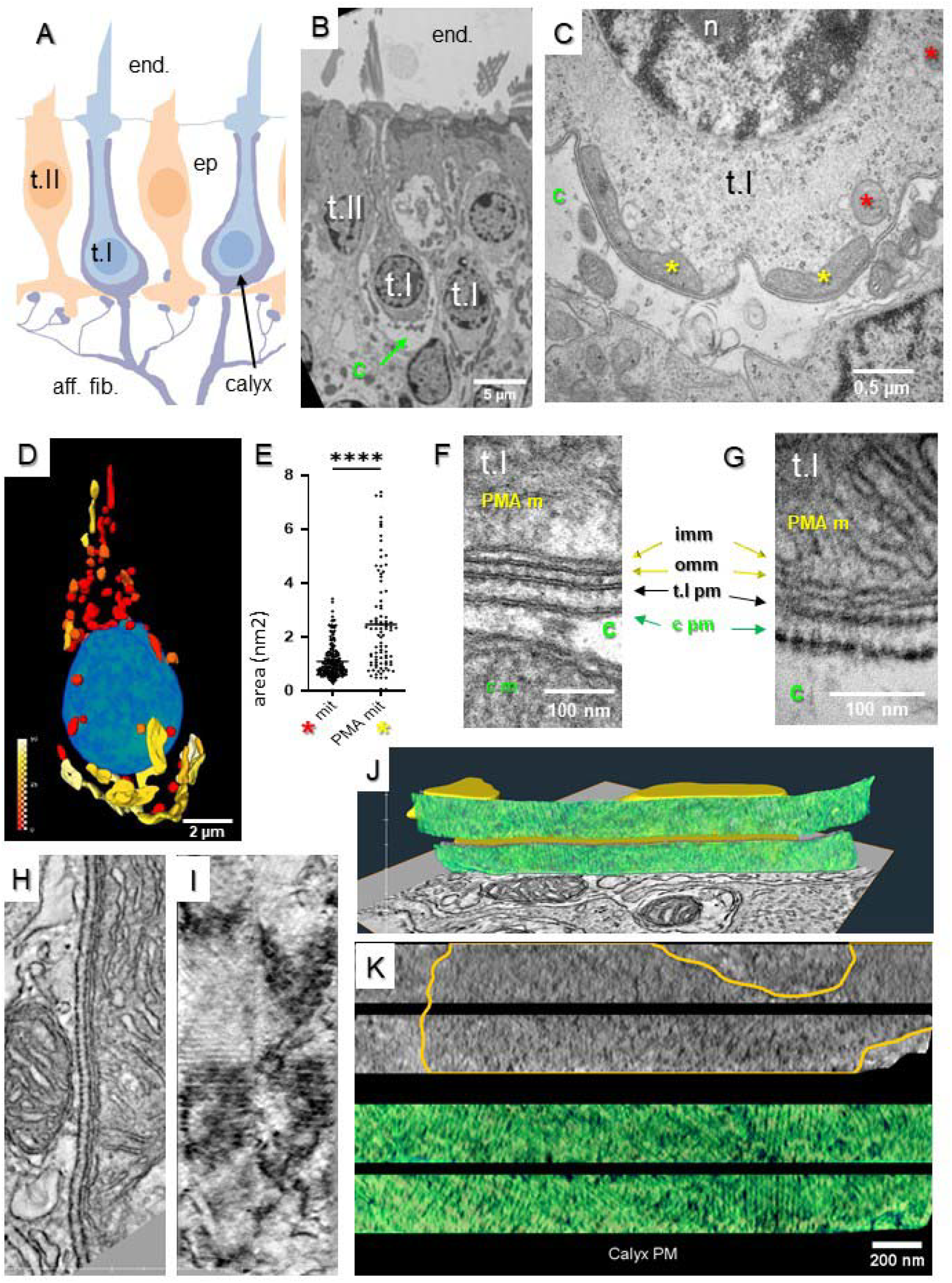
A novel striated junction between the type I hair cell and its calyceal afferent ending in the mouse vestibular sensory organs **A:** Schematic drawing of the utricular epithelium organisation in an adult mouse (t.I, type I cell; t.II, type II cell; calyx; end, endolymph; ep., sensory epithelium; aff. fib., afferent fibres). **B:** Electron micrograph of mouse utricular epithelium showing t.I, t.II and calyx (C). **C:** Adult t.I transverse section, right below the nucleus; yellow *, PMA mitochondria; red *, other mitochondria. **D**: Segmentation from Serial Blockface (SBF) imaging on t.I showing all mitochondria. Blue ovoid: t.I nucleus. The proportion of contact with the plasma membrane for each mitochondrion is represented in colour ranging from no contact (Red) to 50% of surface contact (white-yellow): mitochondria; (see Suppl. Video 1) **E:** Quantification of the size of the mitochondria depending on their proximity to the plasma membrane. We bin the mitochondria in 2 groups, either < 20% or > 20% surface contact; n=325 on 3 segmented SBF cells, unpaired t-test **** p>0,0001. **F-G:** High magnification electron micrographs of striations. Depending on the angle of the section, striations can be visible or not, highlighting the need for electron tomography (imm: inner mitochondrial membrane; omm: outer mitochondrial membrane; m: mitochondria; pm: plasma membrane; t.I: type I HC; c.: calyx.). H-K: ssTEM tomogram **H-I:** transversal image (H) and corresponding in-face view (I) of a tomogram highlighting the periodic striations on the calyx membrane **J:** 3D rendering of serial tomograms segmentation of the calyx membrane (green) and the PMA mitochondria (blue) (see Suppl. Video 2). **K:** pseudo-coloured plasma membranes from segmentations in two serial tomograms (grey: type I HC plasma membrane, green: calyx plasma membrane, orange contour: the surface of contact between the mitochondria and the type I cell plasma membrane).

In the type I HC, as in other cochlear or vestibular sensory HCs, mitochondria are distributed throughout the whole cytoplasm but also form two distinct clusters with likely different physiological functions. One cluster sits just below the cuticular plate, a large apical structure into which the mechanosensory stereocilia implant, and is therefore well-positioned to support mechanoelectrical transduction. Another cluster is found at the basal synaptic pole, well-positioned to sustain neurotransmission (Lysakowski et al., 2022). With transmission electron microscopy (TEM) on ultrathin sections of this synaptic cluster, we found that several mitochondria closely abut the plasma membrane (Figure 1C; adult mouse utricle). These plasma membrane-apposed (PMA) mitochondria were found specifically in the type I HC, throughout the utricular epithelium, in both physiologically and morphologically distinct central (‘striolar’) and peripheral (‘extrastriolar’) zones, and were also noted in the crista ampullaris (Supplemental Figure 1). To our knowledge, PMA mitochondria have not been described in type I HCs in other species.

Serial block-face (SBF) scanning electron microscopy of whole type I HCs in mouse utricles revealed the localisation, size and proportions of the PMA mitochondria (Figure 1D). We confirmed that these mitochondria dominated the basal half of the cell below the cell nucleus, and that they contacted 10-20% of the type I HC plasma membrane at this level. The increased contact with the plasma membrane was correlated with increased size and more elongated shape, as if the mitochondria were spreading along the plasma membrane (Figures 1D-E). Using TEM, we determined that the PMA mitochondria are elongated and observed a remarkable alignment of 4 membranes, the plasma membranes of the calyx and the type I HC, together with the inner and outer membranes of the PMA mitochondria (Figures 1F-H). This revealed two striking features: 1) the outer membrane of each mitochondrion is very closely apposed to the type I HC plasma membrane, with only a 10-15 nm gap between the two structures (Figures 1F-H); and 2) alternating light and dark electron-dense stripes are visible on the plasma membranes of both the type I HC and the post-synaptic calyx (Figure 1G). We refer to these specialised zones as “striated contacts”. Using serial section Electron Tomography (ssET), we confirmed the presence of striations on type I HC and on the facing calyx plasma membranes (Figures 1G,H). We could not find an example of an apposed mitochondrion without colocalised striations (n=5 type I HC). These observations suggest that there may be a novel specific contact between the calyx and the type I HC mediated by close association with a mitochondrion.

Using TEM, we observed that PMA mitochondria in type I HCs first appear relatively late during mouse development, around weaning. 10 % of type I HCs had at least one PMA mitochondrion at P10-14 (n=39 HC), with 24 % at P18 (n=17), 84 % at P25 (n=37), and 85 % in adults >P42 (n=12) (Table S1). During development, the calyx is growing and encapsulating the type I cells from the base to the apex and appears to be fully mature from P14 (Warchol et al., 2019). This suggests that PMA mitochondria appear after the full development of the calyx. It is interesting to note that the appearance of large mitochondria in the basal pole of type I HCs has recently been described in adult mouse (Michanski et al., 2023).

In the adult, ribbons at synapses become sparse in mouse type I HCs (Favre and Sans, 1978; Warchol et al., 2019) while floating ribbons become predominant (Michanski et al., 2023). It was recently reported that NQT becomes a dominant form of synaptic activity in type I HCs as mice mature, while both quantal and non-quantal forms are common in developing animals (Eatock, 2018; Govindaraju et al., 2023; Songer and Eatock, 2013). The concomitant appearance of striated junctions facing PMA mitochondria with NQT synaptic maturation raises the possibility that these structures play a role in NQT. Interestingly, PMA mitochondria are confined to the basal region below the nucleus, while the mature calyx typically extends to the cuticular plate. Given that calyx height determines the magnitude and speed of NQT (Govindaraju et al., 2023), the basal localisation of PMA mitochondria might modulate NQT by regulating local ions or energy availability. Over longer timescales, calyx height could set the overall magnitude and speed of NQT, while PMA mitochondria might fine-tune these processes on shorter timescales.

How do PMA mitochondria in type I HCs compare to other mitochondria-plasma membrane contact sites? These contacts typically vary in their distance from the plasma membrane, ranging from 100 nm to as close as 20 nm in gap junction plates (Cetin-Ferra et al., 2023). Remarkably, only two examples in the literature—bat follicular cells and mouse photoreceptors—show a similarly narrow 10 nm gap (Montes de Oca Balderas, 2021). In both cases, in contrast to type I HCs, the mitochondria are symmetrically mirrored by apposed mitochondria in adjacent cells, a configuration proposed to be linked to cellular synchronisation. Further research is needed to uncover the structural and molecular components of these novel striated contacts and the molecular mechanisms underlying mitochondria tethering in type I HCs. Understanding their role will provide valuable insights into synaptic transmission and broader cellular communication processes.

## Supporting information

Supplementary Video 1_SBF

Supplementary Video 2_sTEM

## Acknowledgements

We thank Ed Parker for sample preparation at the University of Washington supported by Core Grant for Vision Research NEI P30EY001730. The electron microscopy experiments were performed on the PiCSL-FBI core facility, a member of the France-BioImaging national research infrastructure (ANR-10-INBS-04) and a member of the Marseille Imaging Institute, an Excellence Initiative of Aix Marseille University A*MIDEX, a French “Investissements d’Avenir” programme (AMX 19 IET 002). This work was funded by grants from NIH R01 DC013771 to Stone, JS, and from France BioImaging (ANR-10-INBS-04) to N.B. and R.P.

## Supplementary Information

**Supplementary Figure 1.**
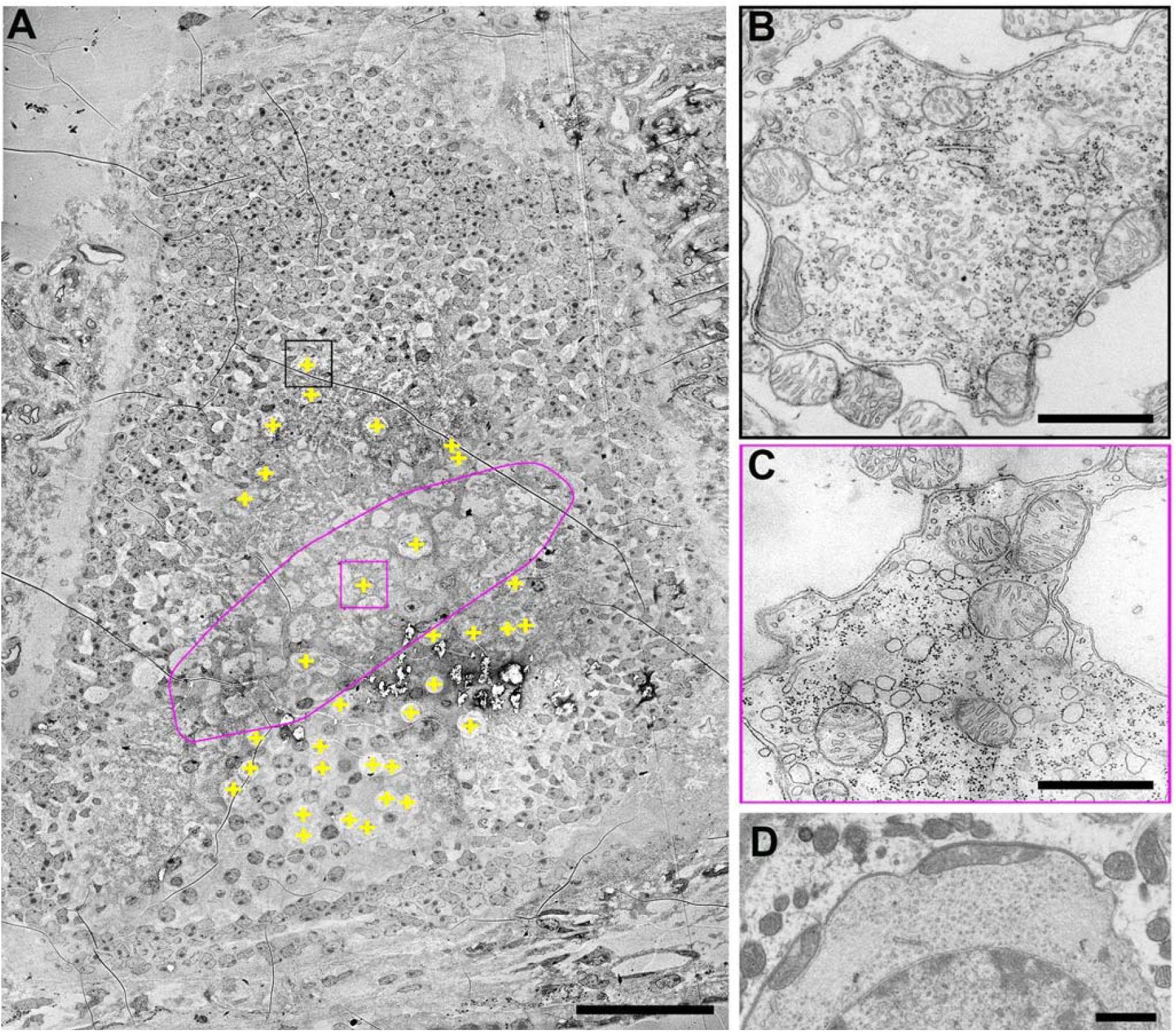
PMA mitochondria were observed in different zones of the utricle - the striola and extrastriola - and in the posterior cristae ampullaris A-TEM overview of the whole utricle. The area outlined in magenta corresponds to the striola, identified by the numerous multicalices. The ‘+’ symbols represent type I cells with at least one PMA. Scale bar, 50 µm. B-D Examples of PMA mitochondria in the extrastriola (B) the striola (C), and the cristae ampullaris (D), scale bar, 1 µm.

**Supplementary Table 1:**
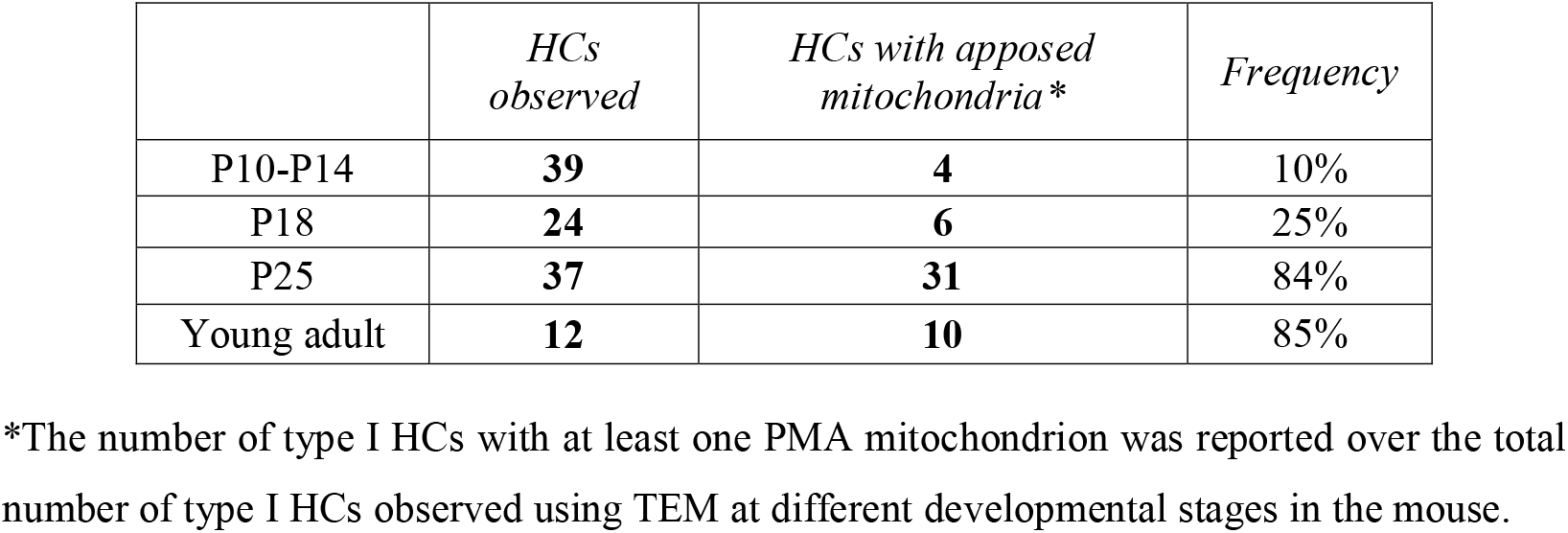
PMA mitochondria become dominant at P25 in type I HC.

|  | *HCs*<br>*observed* | *HCs with apposed*<br>*mitochondria\** | *Frequency* |
| --- | --- | --- | --- |
| P10-P14 | **39** | **4** | 10% |
| P18 | **24** | **6** | 25% |
| P25 | **37** | **31** | 84% |
| Young adult | **12** | **10** | 85% |

\*The number of type I HCs with at least one PMA mitochondrion was reported over the total number of type I HCs observed using TEM at different developmental stages in the mouse.

**Supplementary Video 1:** SBF of a type I HC revealing (PMA) mitochondria in yellow and and non-apposed mitochondria in red, nucleus in bleu.

**Supplementary Video 2:** Serial tomography of a striated junction with a mitochondrion in blue and the plasma membrane in green.

## Material and Methods

### Animals

All mice were C57BL/6J background. Both genders were used in all studies. The University of Washington School of Medicine (Seattle, WA) conducted all animal procedures using approved protocols from their animal care and use committee that conform to the standards of the American Veterinary Medical Association and the National Institutes of Health.

### Electron Microscopy - Serial Block Face SEM sample preparation

Utricles were fixed in 4% glutaraldehyde in 0.1M sodium cacodylate buffer then post-fixed in osmium ferrocyanide (2% osmium, 1.5% potassium ferrocyanide, in 0.1M cacodylate buffer). Then utricles were incubated in 1% thiocarbohydrazide for 20 minutes, then in 2% osmium tetroxide for 30 minutes, then in 1% uranyl acetate overnight in the refrigerator. Next, utricles were stained using Walton’s lead aspartate for 30 minutes at 60°C, then dehydrated in ice cold 30%, 50%, 70%, and 95% ethanol, then 2 changes of 100% ethanol and two changes of propylene oxide. Rinses were performed at each step, and every step was performed at room temperature unless noted. Utricles were embedded in Durcupan resin. The block was attached to the mounting pin using conductive epoxy made by Circuitworks and supplied by SPI. The block was trimmed to ∼ 750 µm on each side and coated with gold using a sputter coater.

### Electron Microscopy - Serial Block Face SEM Acquisition and segmentation

Image stack acquisition was performed on a Teneo VS (ThermoFisher Scientific) in LoVac (30 Pa of water vapour) in ThruSight mode with an acquisition voxel size of 10x10x20 nm and sectioning every 40 nm. The data set was then binned twice in x,y to end up with isotropic voxels of 20x20x20 nm.

The extracellular environment was outlined using the ilastik carving mode (Berg et al., 2019) to generate a first dataset in order to later mask out all the mitochondria that don’t belong to the Type I HC. All mitochondria of the original dataset were segmented using the pixel classification mode of Ilastik using seeds in the Type I HC mitochondria. In parallel, the same original data set was treated with the pretrained CebraNet model to reveal all cellular membranes. Finally combining the three resulting datasets, we could isolate the Type I HC mitochondria and run their morphological analysis in Amira. The nucleus was segmented using the ilastik carving mode. Beyond classical 3D morphometrics (including area and Feret diameters), we measured the proportion of contact between every Type I HC mitochondrion and the plasma membrane by dividing half of the Area of the interaction plate between the mitochondria and the plasma membrane by the Area of the same mitochondria. Movies were generated in Amira and the colour coding of each mitochondrion corresponds to its proportion of contact with the Type I HC plasma membrane which can span from 0% (no contact, red) to about 50% (fully apposed mitochondria, white).

### Electron Microscopy - Serial Tomography sample preparation

Utricles and posterior crista ampullae were prepared as described in (Warchol et al., 2019). Briefly, temporal bones were isolated and fixed in 2.5% glutaraldehyde in 0.1M cacodylate buffer, rinsed and post-fixed with 2% osmium tetroxide in cacodylate buffer. The end organs were dissected and embedded in Eponate (Ted Pella Inc. #18010). Transverse ultrathin (80-90 nm) sections were taken through the extrastriolar/peripheral and striolar/central zones of the organs and were imaged using a Tecnai G2 microscope (FEI) transmission electron microscope equipped with an Veleta camera (Olympus, Japan) digital camera.

### Electron Microscopy - Serial Tomography acquisition and segmentation

350 nm semithin sections were performed using a Leica UC7 ultramicrotome and post-stained with 2% uranyl acetate and Reynolds’ lead citrate. Images were taken with a Tecnai G2 microscope (FEI) at 200 kV. For tomography acquisitions, tilted images (+60°/–60° according to a Saxton scheme) were acquired using Xplorer 3D (FEI) with a Veleta camera (Olympus, Japan). Tilted series alignment and tomography reconstruction were performed using IMOD (Mastronarde and Held, 2017). The segmentation and movie generation was carried out in Amira. To segment the uneven curvature of the plasma membrane, we regularly placed landmarks to generate a surface that was dilated and used as a mask to reveal its striation in an *en face* view.

